# What happens after a blood meal? A transcriptome response from the main tissues involved in egg production in *Rhodnius prolixus*, an insect vector of Chagas disease

**DOI:** 10.1101/2020.06.26.173195

**Authors:** Jimena Leyria, Ian Orchard, Angela B. Lange

**Author notes:** Corresponding author (JL).

## Abstract

The blood-sucking hemipteran *Rhodnius prolixus* is a vector of Chagas disease, one of the most neglected tropical diseases affecting several million people, mostly in Latin America. The blood meal is an event with a high epidemiological impact since in adult mated females it initiates the production of hundreds of eggs. By means of RNA-Sequencing (RNA-Seq) we have examined how a blood meal influences mRNA expression in the central nervous system (CNS), fat body and ovaries in order to promote egg production, focusing on tissue-specific responses under controlled nutritional conditions. We illustrate the cross talk between reproduction and a) lipids, proteins and trehalose metabolism, b) neuropeptide and neurohormonal signaling, and c) the immune system. Overall, our molecular evaluation confirms and supports previous studies and provides an invaluable molecular resource for future investigations on different tissues involved in successful reproductive events. Analyses like this can be used to increase the chances of developing novel strategies of vector population control by translational research, with less impact on the environment and more specificity for a particular organism.

**Author summary:** The blood-sucking hemipteran *Rhodnius prolixus* is one of the main vectors of Chagas disease. The blood meal is an event with a high epidemiological impact since in adult mated females, blood-gorging leads to the production of hundreds of eggs. This work describes an in-depth central nervous system (CNS), ovary and fat body transcriptome analysis, focusing on transcripts related to blood intake which may be relevant in promoting egg production. To date, the principle focus in Chagas disease prevention is on the elimination of triatomine vectors and their progeny. This work will serve as a starting point for initiating novel investigations on targets identified with a potential for use in vector control; for example using specific genes to generated symbiont-mediated RNAi, a powerful technology which provides a novel means in biocontrol against tropical disease vectors.

## Introduction

Insects, which represent more than half of all living organisms on earth, have a close relationship with human beings. To many of them, we can ascribe a negative interaction, for example those that act as carriers of disease. Chagas disease, one of the most neglected tropical diseases, is caused by the protozoan *Trypanosoma cruzi*, which is transmitted to mammalian hosts primarily by blood-feeding insects, the triatomines [1]. This disease affects 6-7 million people, mostly in Latin America, but because of migration the disease has spread to other continents [2]. To date, treatment of the chronic phase of this disease is limited [3], resulting in 2000 deaths per year [1], although it is known that Chagas disease is an under-reported cause of death [4]. The principle focus in Chagas disease prevention is on the elimination of triatomine vectors from human homes. Currently, the most heavily used option is chemical control, although resistance to these insecticides has been reported in the last decade [5]. Furthermore, the devastating impact of chemical insecticides on the environment and other organisms, such as beneficial insects, can no longer be ignored [6].

Triatomines have developed an integrated control over the reproductive system, whereby different tissues work with extreme precision and coordination to achieve successful production of progeny. There are three tissues that work in concert to promote reproduction; the central nervous system (CNS), fat body and ovaries. The CNS contains neuroendocrine cells that synthesize neuropeptides involved in the coordination of events that promote egg production. These neuropeptides are produced as large precursors, which are then cleaved and modified to become biologically active neuropeptides [7]. These neuropeptides are secreted as neuromodulators or neurohormones to act via specific receptors [8]. With regard to reproduction, these receptors are located on the fat body and ovaries. The fat body is a multifunctional organ analogous to vertebrate adipose tissue and liver. It is considered an interchanging center, remotely integrating with the CNS to regulate physiology by sensing hormonal and nutritional signals and responding by mobilizing stored nutrients such as proteins, carbohydrates and lipids, for use in egg formation, or during periods of inactivity or nutritional shortage [9-10]. Apart from these storage functions, the fat body is also involved in the regulation of hematopoiesis, innate immune homeostasis and detoxification [10].

In oviparous organisms, including triatomines, embryonic development occurs apart from the maternal body. Egg survival, therefore, depends on the utilization of previously stored material taken up by the oocytes, such as proteins, lipids, carbohydrates and other minor components, all of which are synthesized mainly by the fat body [11]. Insect oocytes are specific structures designed to select, internalize, and store nutrients, such as yolk granules and lipid droplets. The process of yolk deposition is termed vitellogenesis, which represents a phase of accelerated egg growth leading to the production of mature eggs in a relatively short period of time [11]. The CNS-fat body-ovary axis is essential for triatomines to produce viable eggs. Interestingly, the trigger for this interaction is a single blood meal. Although in some colonies of the triatomine, *Rhodnius prolixus*, unfed females can make a small number of eggs from resources that may remain after moulting to an adult (autogeny) [12], the large batch of eggs is triggered by ingestion of a blood meal. After a blood meal, a *R. prolixus* female can produces 30–50 eggs during the following three weeks [13]. For this reason, knowledge of the molecular and cellular mechanisms used in egg formation are essential to develop novel strategies of vector population control.

In addition to being a main vector of Chagas disease, with high epidemiological relevance for easily colonizing domestic habitats [14], *R. prolixus* has been the subject of intense investigations over the past century, which have contributed to our understanding of important aspects of metabolism and physiology in insects [15]. It is important to highlight that the complete genome of *R. prolixus* has been published [16] and, therefore, many new questions can be asked and answered with regard to insect physiology/endocrinology. Next-generation sequencing allows us to study biological systems at the genomic level to link mRNA sequences with specific biological functions of specific tissues during a particular stage or state. Recently, by transcriptome analysis we reported an up-regulation of transcripts involved in insulin-like peptide/target of rapamycin (ILP/ToR) signaling in unfed insects. However, we demonstrated that this signaling pathway is only activated in the fat body and ovaries of fed insects. Thus, we demonstrated that unfed females are in a sensitized state to respond to an increase of ILP levels by rapidly activating ILP/ToR signaling after a blood meal [17]. Here, we examine how a blood meal influences CNS, fat body and ovary gene expression to promote egg production; focusing on details associated with tissue-specific responses in particular nutritional states. Our data opens up avenues of translational research that could generate novel strategies of vector population control, with less impact in the environment and with more specificity for a particular organism, such as using specific genes to generated symbiont-mediated RNAi; a powerful technology which provides a potential means in biocontrol against tropical disease vectors [18].

## Materials and Methods

### Insects

Insects were maintain in incubators at 25°C under high humidity (∼50%). Newly-emerged adult females (10 days post-ecdysis) were segregated, and placed together with a recently fed male to copulate. Then, female insects were fed on defibrinated rabbit blood (Cedarlane, Burlington, CA) to initiate egg growth. Only insects that fed 2.5 to 3 times their initial body weight were used for the experiments. CNS, fat body (FB) and ovaries (OV) from adult mated females were dissected at 10 days post-ecdysis for the unfed condition (UFC) and 3 days post-feeding as the fed condition (FC), according to Leyria et al. [17]. Insects in the fed condition will have begun vitellogenesis and egg growth.

### Transcriptomic data analysis

Read sequences were obtained from Leyria et al. [17]. This study reported transcriptomes of CNS, FB and OV from fed and unfed females. The raw sequence dataset of this project is registered with the National Center for Biotechnology Information (NCBI) under PRJNA624187 and PRJNA624904 bioprojects. A detailed description of our bioinformatic pipeline can be found in Leyria et al. [17]. Briefly, CNS, OV and ventral and dorsal FB of *R. prolixus* females were dissected in cold autoclaved phosphate buffered saline (PBS, 6.6 mM Na_2_HPO_4_/KH_2_PO_4_, 150 mM NaCl, pH 7.4). Three independent experiments were analyzed (n = 3) with each n composed of a pool of 10 tissues. RNA extraction was performed with Trizol reagent (Invitrogen by Thermo Fisher Scientific, MA, USA), followed by DNase treatment (Millipore-Sigma, WI, USA) and then repurified with PureLink RNA Mini Kit (Ambion by Thermo Fisher Scientific, MA, USA). Libraries for sequencing were made from high quality RNA that were generated using NEBNext Ultra RNA Library Prep Kit for Illumina (New England Biolabs, MA, USA) following manufacturer’s recommendations. The libraries were sequenced on Illumina HiSeq platforms (*HiSeq 2500*) at the Novogene sequencing facility (California, USA). Raw data were recorded in a FASTQ file, which contains sequence (reads) and corresponding sequencing quality information. Fastq format were first processed through in-house perl scripts, where clean data (clean reads) were obtained by removing reads containing the adapter, reads containing ploy-N and low quality reads from raw data. Also, Q20, Q30 and GC content from the clean data were calculated. All the downstream analyses were based on the clean data [17].

### Differential expression analysis

First, clean reads were aligned to the reference genome using HISAT2 software. After that, HTSeq v0.6.1 was used to count the number of reads mapped to each gene. FPKM (expected number of Fragments Per Kilobase of transcript sequence per Millions base pairs sequenced) of each gene were calculated based on the length of the gene and numbere of reads mapped to the gene. In general, an FPKM value of 0.1 was set as the threshold for determining whether the gene is expressed or not. Differential expression analysis of two nutritional conditions were performed using the DESeq R package (1.18.0). DESeq provides statistical routines for determining differential expression in digital gene expression data using a model based on the negative binomial distribution. The resulting *P*-values were adjusted using the Benjamini and Hochberg’s approach for controlling the false discovery rate. Genes with an adjusted *P*-value < 0.05 found by DESeq were assigned as differentially expressed. Gene Ontology (GO) enrichment analysis of differentially expressed genes was implemented by the GOseq R package, in which gene length bias was corrected. GO terms with corrected *P*-value less than 0.05 were considered significantly enriched by differential expressed genes.

### Validation of quantitative gene expression

To validate Illumina sequencing results, 7 genes were chosen at random and their expressions were analyzed by quantitative polymerase-chain reaction (RT-qPCR) [17]. Briefly total RNA was extracted as described above, followed by cDNA synthesis using the High Capacity cDNA Reverse Transcription Kit (Applied-Biosystems, by Fisher Scientific, ON, Canada). RT-qPCR was performed using an advanced master mix with super green low rox reagent (WISENT Bioproducts Inc, QC, Canada). Three independent experiment were performed (n=3) with each n composed of a pool of 5 tissues. Each reaction contained 3 technical replicates and were carried out using a CFX384 TouchTM Real-Time PCR Detection System (BioRad Laboratories Ltd., Mississauga, ON, Canada). The primers used (by Sigma-Aldrich, ON, Canada) for amplification are shown in S1 Table. Quantitative validation was analyzed by the 2^^-ΔΔCt^ method. All reactions showed an amplification efficiency higher than 95 %. β-actin, which was previously validated for transcript expression in FB and OV from *R. prolixus* at different nutritional conditions [17], was used as the reference gene. For each pair of primers a dissociation curve with a single peak was seen, indicating that a single cDNA product was amplified. Specific target amplification was confirmed by automatic sequencing (Macrogen, NY, USA). The correlation coefficient between Illumina RNA sequencing and RT-qPCR data was analyzed by the Pearson test.

### Lipid and carbohydrate measurements

Ovaries and ventral and dorsal FB were dissected from insects during UFC and FC under cold *R. prolixus* saline (NaCl 150 mM, KCl 8.6 mM, CaCl_2_ 2.0 mM, MgCl_2_ 8.5 mM, NaHCO_3_ 4.0 mM, HEPES 5.0 mM, pH 7.0). Total lipids and carbohydrates from tissues were measured by colorimetric assays as previously described [19]. Briefly, the tissues were placed in either 500 μl of isopropanol (for lipid quantification) or 500 μl 10 % cold trichloroacetic acid (TCA, for carbohydrate quantification), homogenized and then centrifuged for 10 min at 20 °C and 8000 g. For lipid quantification, 400 μl of the supernatants were transferred to 1.5 ml tubes containing 100 μL of 1 M KOH. Then, the tubes were incubated at 60°C for 10 min and once they were cool, 100 μl of sodium periodate solution (11.6 mM sodium periodate in 2 N glacial acetic acid) was added. After 10 min of incubation at room temperature, 600 μl of chromogenic solution (40 ml of 2 M ammonium acetate, 40 ml isopropanol, 150 ml acetyl acetone) were added to the samples and incubated for 30 min at 60 °C. The resultant colour was measured at 410 nm using a plate reader spectrophotometer (Cytation 3 Imaging Reader, BioTek, Winooski, VT, USA). A standard curve of triglycerides ranging from 0 to 60 μg was run independently and in parallel with the experimental samples. FB and OV carbohydrate content was measured using the anthrone colorimetric assay. Briefly, 50 μl of the supernatants after TCA precipitation were mixed with 500 μl of anthrone solution (26 mM anthrone, 1.31 mM thiourea, 66 % sulfuric acid) and incubated for 20 min at 100 °C. The samples were allowed to cool in the dark for 15 min and then quantified at 620 nm using a plate reader spectrophotometer described. A standard curve was run in parallel with the experimental samples using a 0 - 40 μg range of trehalose. Proteins were measured according to Bradford [20] processing the tissue as described previously [17]. Three independent experiments were analyzed (n=3) for each measurement with each n composed of a pool of 5 tissues.

## Results and Discussion

We were surprised to observe no major gene differences in the CNS between the UFC and FC. None of the GO functional terms were enriched in the CNS under these different nutritional states. We chose 3 days post-blood meal as the fed condition because of the morphological changes observed in the FB and OV (Leyria et al., 2020). The days chosen to monitor transcriptional regulation are appropriate for FB and OV but apparently not for CNS. Possibly for the CNS, transcriptional regulation begins early after a blood meal to control remotely molecular, biochemical and physiological changes that we then observed in the FB and OV during the FC. Using *R. prolixus* adult insects, Sterkel et al. [21] reported a quantitative proteomic analysis of the post-feeding response from CNS in 3 different conditions: unfed, 4 h and 24 h after blood intake. These researchers found only 4 neuropeptides (NVP-like, ITG-like, kinin-precursor peptide and neuropeptide-like precursor 1 (NPLP1)) that were significantly up-regulated in response to the blood meal. Taken together, this appears to indicate that the changes in the transcriptional and protein levels in the CNS of *R. prolixus* adult insects occur quickly or more slowly so that it is difficult to find any changes. For this reason, below, we focused our attention on the FB and OV and reflect on the CNS transcriptome analysis when making reference to peptide/hormone signaling.

To validate Illumina sequencing, 7 mRNAs were chosen and their relative transcript abundance in FB and OV in both nutritional states monitored by RT-qPCR. A good correlation was found between RNAseq and RT-qPCR data; the Pearson tests were 0.9311 (to FB) and 0.9109 (to OV), with a statistical significance of p<0.01. Multiples genes from these transcriptomes were also validated using RT-qPCR by Leyria et al. [17].

### GO enrichment analysis

Nutrients are essential for energy homeostasis of any organism and important changes in nutrient stores occur between feeding and non-feeding periods and also more remarkably in adult insects during reproductive processes [9, 22]. GO enrichment was used to assign a functional classification to differentially expressed genes (DEGs). All DEGs categorize into two main groups: cellular components and biological processes. In cellular components, they are divided into 21 terms which are significantly up-regulated in FB_FC with respect to FB_UFC (Fig 1A). The most represented cellular components terms are cell parts involved in protein synthesis, as is to be expected since the FB is the main synthesis and secretory organ responsible for the production of virtually all haemolymph proteins. With regard to biological processes, the main terms in the FB are involved in biosynthesis and lipid and carbohydrate metabolism (Fig 1B). Recently, by examining KEGG enrichment we reported that the “ABC transporters pathway”, transporters which use energy to translocate substrates (e.g., sugar, lipid and peptides) across cell membranes, is up-regulated in FB_FC, which shows that the synthesized nutrients are released, in this case likely to support vitellogenesis [17]. In the OV, the main terms of cellular components and biological processes which are significantly up-regulated in OV_FC with respect to OV_UFC are related to lipid, carbohydrate and protein metabolism, insect hormone biosynthesis, and yolk granule formation (specialized structures which stores all nutrients which are used as substrates for embryogenesis and maintenance of the newly hatched nymph) (Fig 2A and B). These nutrients are mostly proteins, lipids and carbohydrates, produced by the FB, released into the hemolymph and subsequently taken up by the oocytes [23]. As we anticipated in light of the results of the GO enrichment, lipid, protein and carbohydrate levels in the FB and OV are increased in fed females (Fig 3A and B), as reported in other vectors of Chagas’ disease [22, 24-25]. In addition, it is clear that stored proteins are always the major component in both tissues, followed by lipids and then carbohydrate stores.

**Fig 1.**
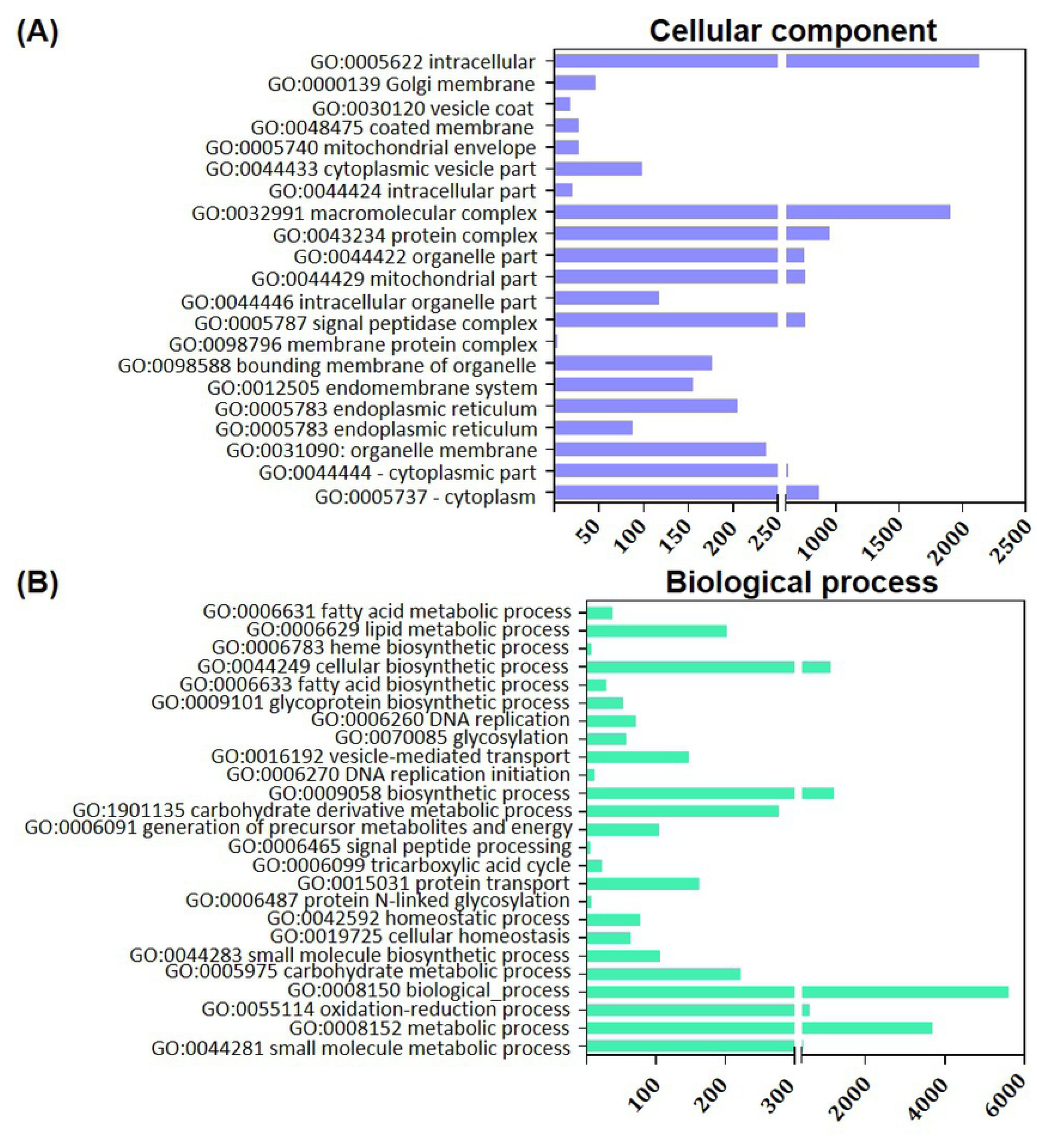
Distribution of differentially expressed genes (DEG) in the fat body annotated by GO enrichment analysis, associated with cellular components and biological processes. The GO enrichment bar chart presents the number of DEG enriched in cellular component (**A**) and biological process (**B**). The y-axis is GO terms enriched and the x-axis is the number of DEG. GO terms with corrected *P*-value less than 0.05 were considered significantly enriched in DEG (comparing FB_FC vs FB_UFC). The most significant enriched terms were selected. FB_FC, fat body in fed condition; FB_UFC, fat body in unfed condition.

**Fig 2.**
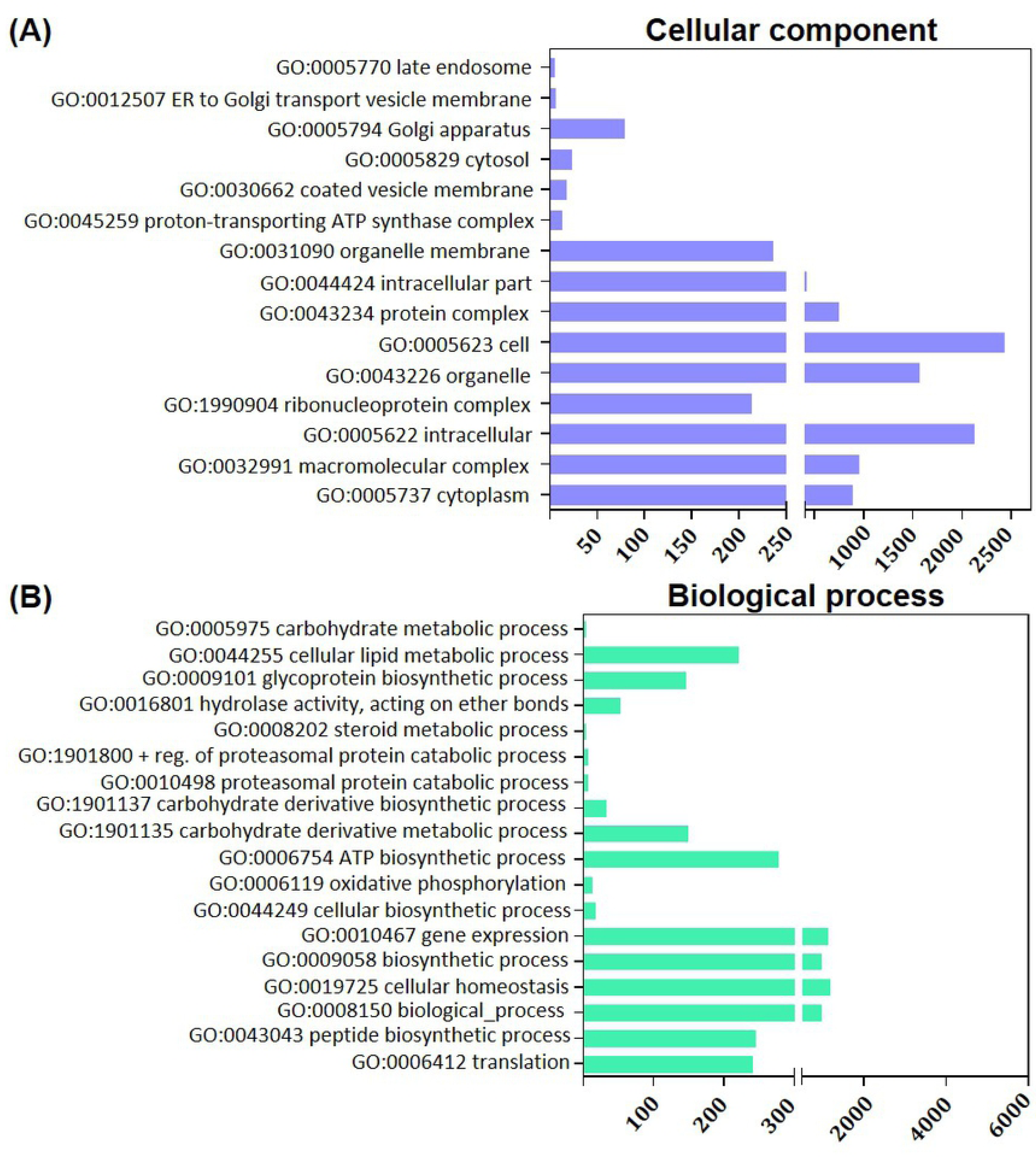
Distribution of differentially expressed genes (DEG) in ovaries annotated by GO enrichment analysis, associated with cellular components and biological processes. The GO enrichment bar chart presents the number of DEG enriched in cellular component (**A**) and biological process (**B**). The y-axis is GO terms enriched and the x-axis is the number of DEG. GO terms with corrected *P*-value less than 0.05 were considered significantly enriched in DEG (comparing OV_FC vs OV_UFC). The most significant enriched terms were selected. OV_FC, ovary in fed condition; OV_UFC, ovary in unfed condition.

**Fig 3.**
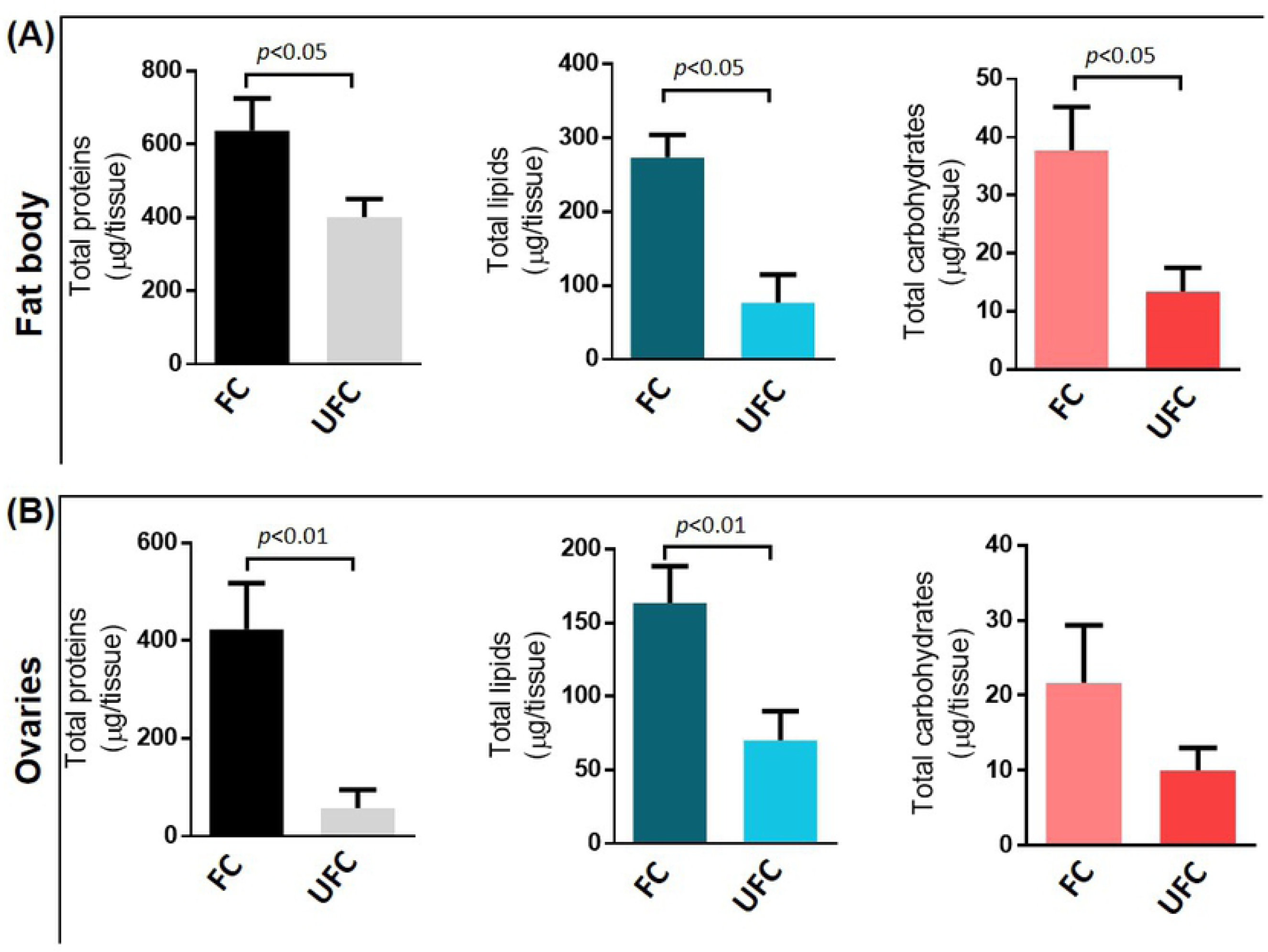
Protein, lipid and carbohydrate content in the fat body and ovaries of females during the unfed (UFC) and fed condition (FC). Fat body and ovaries were dissected from adult females during the unfed and fed condition. The tissues were homogenized and nutrient extracted and quantified as described in Materials and Methods. The results for total lipid, carbohydrate and protein content were graphed as the mean ± Standard Error of the Mean (SEM) from three independent experiments. Graphs and statistical tests were performed using GraphPad Prism 7 (GraphPad Software, CA, USA, www.graphpad.com). All datasets passed normality and homoscedasticity tests. The statistical significance of the data was calculated using Student’s t-test. A P value < 0.05 was considered statistically significant.

### Protein and hormone analysis

Vitellogenins (Vgs), the main yolk protein precursors (YPPs), are large molecules synthesized predominantly by the FB, secreted into the hemolymph and then transported to the OVs. The number of genes encoding insect Vgs varies from one to several depending on the species [26]. Our results show that the mRNA levels for Vgs are considerable higher in the FB with respect to the OV, which is not surprising (Fig 4A). In the FB transcripts for *Vg1* and *Vg2* are up-regulated during vitellogenesis (FB_FC), with Vg1 having the highest expression (Fig 4A and S2 Table). In *Triatoma infestans*, a triatomine related to *R. prolixus*, Vg1 and Vg2 genes are expressed at relatively low levels during the UFC and both Vg transcripts are up-regulated after blood-feeding [27]. Recently, by KEGG enrichment we reported “amino sugar and nucleotide sugar metabolism” and “N-Glycan biosynthesis” are pathways up-regulated in the FB of fed females [17]. Glycosylation is a critical post-translational modification to obtain the proper protein structure for adequate protein function and for Vgs glycosylation is a step necessary for folding, processing and transport to the oocyte [28]. As previously reported in *R. prolixus* [29], our results suggest that Vg synthesis also occurs in the OV, with *Vg* transcripts up-regulated after a blood meal and *Vg1* levels higher than *Vg2* (Fig 4A and S2 Table). Interestingly, in *T. infestans* the *Vg2* transcript is quantitatively more important that *Vg1* in OVs after feeding [27]. The *vitellogenin receptor* (VgR) mRNA expression, the endocytic receptor responsible for Vg uptake by oocytes, is up-regulated in OV from unfed insects (Fig 4A and S2 Table), contrary to expectation since Vg uptake occurs after a blood meal. However, as expected the main KEGG enrichment pathways involved in receptor-mediated endocytosis signaling (endocytosis, lysosome and phagosome pathways) are enriched in OV_FC of *R. prolixus* [17]. This result indicates that even when the OV expresses high endocytic receptor transcript levels in the UFC, only after a blood meal does the endocytic process occur. Similarly, in cockroaches VgR mRNA levels remain low during the vitellogenic phase [30-31]. A similar pattern of high VgR mRNA levels in non-reproductive stages and low levels during vitellogenesis is found not only in insects but also in oviparous vertebrates [32-33]. It can interpreted as a recycling of VgR protein during the vitellogenetic period. In contrast to these observations, in the mosquito *A. aegypti*, VgR mRNA starts to rise one day after the adult moult, continues to increase dramatically during the vitellogenic period, and then peaks one day after the blood meal [34]. On the other hand, using *R. prolixus* females, Oliveira et al. [35] described another YPP, a 15-kDa protein called *Rhodnius* heme binding protein (RHBP), which works as an antioxidant agent in hemolymph. After the blood meal, a large amount of heme is released from hemoglobin, crosses the digestive barrier and reaches the hemolymph, where it is sequestered by RHBP. Here, we show that in the FB, *RHBP* mRNA levels are up-regulated in females 3 days after feeding (Fig 4A and S2 Table). The increase of synthesis of YPPs in FB_FC coincides with the KEGG analysis reported recently, where we show an enrichment of “biosynthesis of amino acids pathway” [17].

**Fig 4.**
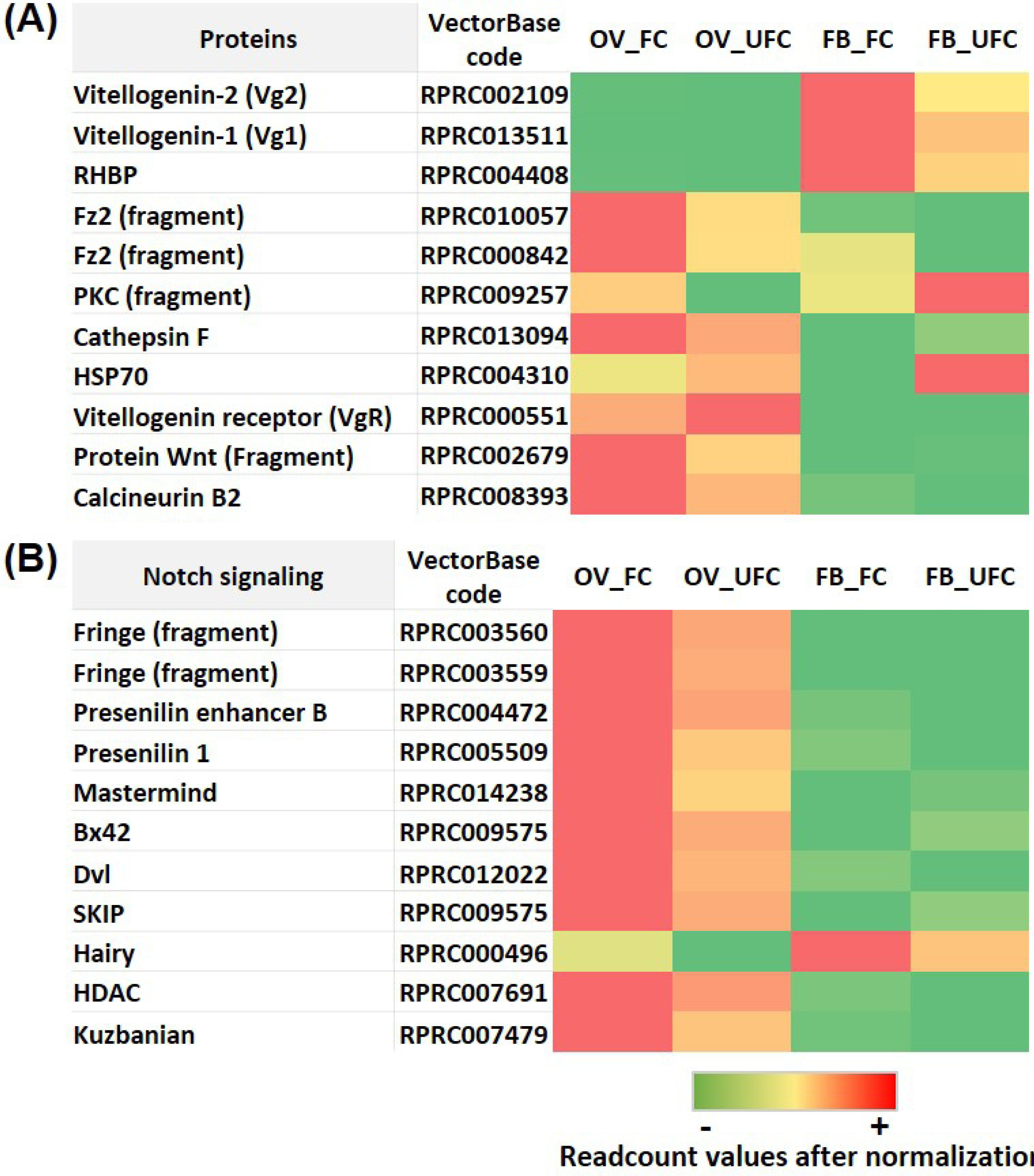
Heat map comparing the mRNA expression levels of proteins related to reproduction (A) and Notch signaling pathway (B) in fat body and ovaries of females in different nutritional conditions. The input data is the readcount value from the gene expression level analysis after normalization and is presented by means of a colour scale, in which green/yellow/red represent lowest/moderate/highest expression. DESeq was used to perform the analysis.

The Wnt signaling pathway was first discovered as a key event in *D. melanogaster* development [36]. The Wnt (glycoprotein ligand) and Frizzled (Fz, transmembrane Wnt receptor) proteins interact with structural components at the cell surface to initiate the signaling cascades that result in transcriptional regulation of gene expression. In *A. aegypti*, a fundamental role of Frizzled 2 (Fz2) was reported in egg production [37]. Here, we find that *Wnt* and *Fz2* mRNA levels are up-regulated in OV_FC (Fig 4A and S2 Table). Additionally, Wnt and ToR signaling interact synergistically in the vitellogenic process [37] and supporting this finding, we showed mToR signaling is active in OV_FC [17]. Also, the non-canonical Wnt pathway indicates that Wnt/Fz signaling leads to the release of intracellular calcium through trimeric G proteins [37]. The calcium release and intracellular accumulation activates several Ca^2+^-sensitive proteins, including protein kinase C (PKC), calcineurin and calcium/calmodulin-dependent kinase II (CamKII). In *A. aegypti* it was found that juvenile hormone (JH) activates the phospholipase C (PLC) pathway and quickly increases the levels of Ca^2+^ for the activation and autophosphorylation of CaMKII, which is involved in patency development [38]. On the other hand, in many animals, a rise in intracellular Ca^2+^ levels is the trigger for egg activation, the process by which an arrested mature oocyte transitions to prepare for embryogenesis. Genetic studies have uncovered essential roles for the calcium-dependent enzyme calcineurin in *Drosophila* egg activation [39]. By DEG analysis, we demonstrate an up-regulation of *PKC* and *calcineurin* in OV from fed insects (Fig 4A and S2 Table).In *R. prolixus*, earlier studies by Ilenchuk et al. [40] suggested that a PKC might be involved in patency and Vg uptake but until now the receptors or molecular mechanisms responsible for this cascade are unknown. The results we observe in vitellogenic oocytes of *R. prolixus* could be indicative of a relationship between patency and Wnt/Fz2/Ca^2+^ signaling. Methoprene-tolerant (Met), which encodes a transcription factor of the bHLH-PAS family, was reported to be a JH receptor [41]. *Krüppel homolog 1* (Kr-h1), identified as the main JH primary-response gene activated by Met [41], is up-regulated in OV_FC (Fig 4A and S2 Table), which supports the hypothesis that in *R. prolixus*, JH is working directly on OVs to stimulate egg formation.

Heat shock proteins represent different protein families based on their sequence homology and molecular masses. Among them, Heat shock protein 70 family (Hsp70) is highly conserved. The expression of Hsp70 is considered a good marker for the inducible stress response in an organism [42]. In *T. infestan*s Hsp70 is strongly expressed in unfed insects [43]. Similarly, in *R. prolixus*, we find that *Hsp70* is up-regulated in the FB from unfed females (Fig 4A and S2 Table), a condition inherently associated with a stressful situation. Glucose-regulated protein of 78 kDa (Grp78) is a member of the Hsp70 family which acts as a chaperone to facilitate protein folding and to inhibit protein aggregation of new peptides. Interestingly, in *Locusta migratoria*, Grp78 was reported as a regulatory factor of Vg synthesis and cell homeostasis in the FB via JH signaling [44]. In *R. prolixus*, we show a significant up-regulation of Grp78-like protein in both FB and OV from fed insects (Fig 4A and S2 Table). This result suggests a novel regulatory mechanism involved in the vitellogenic process of *R. prolixus.*

Notch is a receptor that directly translates information of cell-cell contact to gene expression in the nucleus [45]. By KEGG analysis, it was demonstrated that *Notch signaling* is up-regulated in the OV from fed females [17]. Here, we find that transcripts involved in Notch developmental functions, such as Fridge, presenilin enhancer 2 (PEN-2) and presenilin*-*1, are up-regulated in OV_FC (Fig 4B and S2 Table). Mastermind is an essential nuclear factor that supports the activity of Notch [46]. In OV_FC of *R. prolixus, mastermind* transcriptional factor is up-regulated, as well as Bx42 (Fig 4B and S2 Table), an essential factor which through Notch is involved in the formation of different tissues during embryogenesis [47]. In *Blattellagerm anica*, it was demonstrated that Notch is important in maintaining the proliferative and non-apoptotic state of follicular cells, as well as in differentiation of the posterior follicular cell population [48]. It is likely that the up-regulation of this signaling in *R. prolixus* after a blood meal is related to follicular cell metabolism during egg growth.

During vitellogenesis, JH titres are expected to increase, since JH is one of the main hormones involved in Vg synthesis. In insects, the corpora allata (CA), a pair of endocrine glands associated with the brain, are responsible for the synthesis of this sesquiterpenoid hormone [41]. By KEGG analysis, two pathways related to JH, “Insect hormone biosynthesis” and “Terpenoid backbone biosynthesis”, are up-regulated in the FB and OV during the FC [17]. Here, we find that the levels of enzymes responsible for the synthesis of JH, in general, display an up-regulation in the OV and a non-statistically significant increase in the FC with respect to UFC (Fig 5A and S2 Table). It is important to highlight that *R. prolixus* allatectomized immediately after emergence as adults, continue to make a few eggs [49]. This finding suggests an alternate source for JH apart from the CA, but so far, this source has not been identified. Our results suggest that vitellogenic FB and OVs could be capable of synthesizing JH. In addition, insect cytochrome P450s include a group of different enzymes involved in detoxication and biosynthesis of ecdysteroids and JH [50-51]. Previously, by KEGG analysis, we reported an up-regulation of metabolism of xenobiotics by cytochrome P450 in FB_FC, possibly because of an increase in hormone synthesis or/and a detoxification after a blood meal [17]. Allatostatin-C (ASTC) is a family of peptides originally associated with the control of CA activity but now known to be pleiotropic. ASTC and its paralog, ASTCC, are very similar peptides, likely generated by gene duplication, and their receptors possibly have a common ancestor as well [52]. We find a significant up-regulation of *ASTCC* mRNA expression in OV_UFC (Fig 5A and S2 Table), but so far, there is no evidence about the specific role of this peptide on OVs.

**Fig 5.**
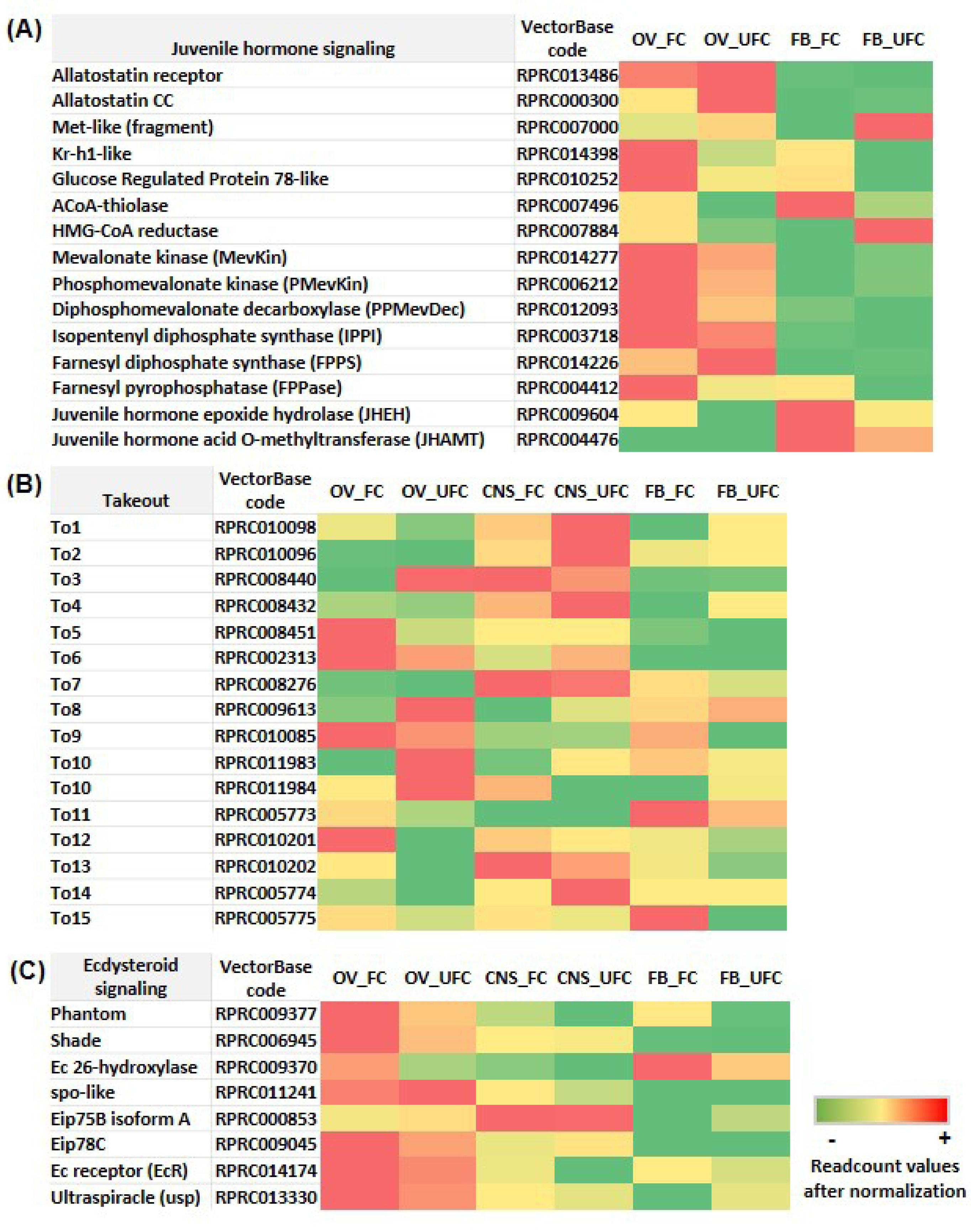
Heat map comparing the mRNA expression levels of molecules involved in juvenile hormone signaling (A), *takeout* genes (B), and ecdysteroid signaling (C) in fat body and ovaries of female adults in different nutritional condition. The input data is the readcount value from the gene expression level analysis after normalization and is presented by means of a colour scale, in which green/yellow/red represent lowest/moderate/highest expression. DESeq was used to perform the analysis

JH is transported from the site of synthesis to target tissues by a haemolymph carrier protein called juvenile hormone-binding protein (JHBP). JHBP protects JH molecules from hydrolysis by esterases present in the insect haemolymph [53]. The takeout genes (*To*) were discovered as a circadian-regulated gene and belong to the JHBPs family [54]. The *To* genes modulate various physiological processes, such as behavioral plasticity in the migratory locust *L. migratoria* and feeding in *D. melanogaster* [55-56]. In the brown planthopper *Nilaparvata lugens*, the *To* family of genes were reported to be regulated by JH signaling [57]. Fifteen such genes were identified in the antenna of *R. prolixus* [58]. Here, we find that *To* genes have a unique pattern of expression according to the tissue analyzed and feeding condition (Fig 5B and S2 Table). *To1, To2, To4* and *To7* mRNA expression is highly expressed in the CNS of unfed insects, suggesting that starvation could induce the expression of these genes. *To9, To11, To12* and *To15* mRNA expression is significantly increased in the FB from females after a blood meal, *To5, T12* and *T13* transcripts show a significantly increased expression in OV_FC (Fig 5B and S2 Table). This is the first report of an analysis of *To* genes in different tissues involved in reproduction in *R. prolixus*, providing new insights into the mechanisms involved in egg formation.

Ecdysteroids are also critical developmental hormones involved in the regulation of molting and metamorphosis. The prothoracic glands (PGs) are the major source of these ecdysteroids in larvae, but PGs are absent from adult insects, where alternative sites of ecdysteroid production have been described. Cardinal-Aucoin et al. [59] reported that in *R. prolixus*, between days 3 and 4 after a blood meal, ovarian ecdysteroid content increased 4–5 fold to a level that was sustained for the duration of egg development. This pattern is similar to that seen in the hemolymph ecdysteroid titer. Two interpretations were proposed a) the ovary passively absorbs hemolymph ecdysteroids or b) the ovary produces the ecdysteroids found in the hemolymph. After a blood meal, we find up-regulation of 3 enzymes involved in ecdysteroid synthesis in the OV, *Shade, Phantom* and *26-hydroxilase*, supporting the second hypothesis (Fig 5C and S2 Table). Similarly, in *Tribolium castaneum*, an insect with the same type of ovaries (telotrophic meroistic), 20-hydroxyecdysone (20E) and its receptors are required for ovarian growth, oocyte maturation and follicle cell differentiation [60]. Overall, this work suggests that the OVs in *R. prolixus* females are an alternate source for ecdysteroid synthesis.

### Carbohydrate analysis

The main blood sugar in insects is trehalose, a sugar that consists of two glycosidically linked glucose units. Trehalose homeostasis is controlled by trehalose-6-phosphate synthase, the main enzyme involved in trehalose synthesis by the FB; trehalose transporter (TRET), which has a particular direction of transport depending on the trehalose gradient, and trehalases, specifically two isoforms, soluble (TRE-1) and membrane-bound (TRE-2), involved in the conversion of trehalose to glucose to generate energy [61-62]. DEG analysis reveals that *trehalose-6-phosphatase synthase* and *TRET* are up-regulated in the FB during the FC (Fig 6 and S2 Table). It is widely accepted that the vitellogenic process is an event with high energy demands. Thus, trehalose synthesis and release by TRET after a blood meal are steps necessary to provide energy to support successful vitellogenesis as well as trehalose to be taken up by developing oocytes, which accumulate carbohydrates as a resource for embryogenesis [63]. Supporting this finding, specific *phospholipase A2-like* mRNA (RPRC008617) is up-regulated in FB_FC (Fig 7 and S2 Table). This belongs to a group of enzymes that are involved in either the formation or release of trehalose from FB cells [64]. In addition, after a blood meal, we find that *trehalose-6-phosphatase synthase* is down-regulated in OV (Fig 6 and S2 Table). Therefore, the trehalose that is stored in the OV to promote glycogen synthesis must be incorporated from extra-ovarian sources. In *R. prolixus*, it has been suggested that TRE-2 in ovaries could react directly with trehalose in the haemolymph supporting the idea that hydrolysis of trehalose at the cellular surface could be an obligatory step to provide glucose for carbohydrate accumulation by oocytes [65]. The researchers found that trehalase activity seemed not to be regulated at the transcriptional level after a blood meal. In addition, here we find that *TRE-2* is up-regulated in OVs but in unfed females (Fig 6 and S2 Table). We hypothesize that glucose obtained by the breakdown of trehalose could participate in the regulation of the energy necessary (contributed by different tissues, including OVs) to maintain overall metabolism of the insect until physiological conditions improve. An interesting finding from our results is that *TRET* is more than 6-fold up-regulated in OVs of fed insects (Fig 6 and S2 Table), supporting the hypothesis that a direct trehalose uptake from the hemolymph by TRET is the most important process involved in the storage of carbohydrates in ovaries.

**Fig 6.**
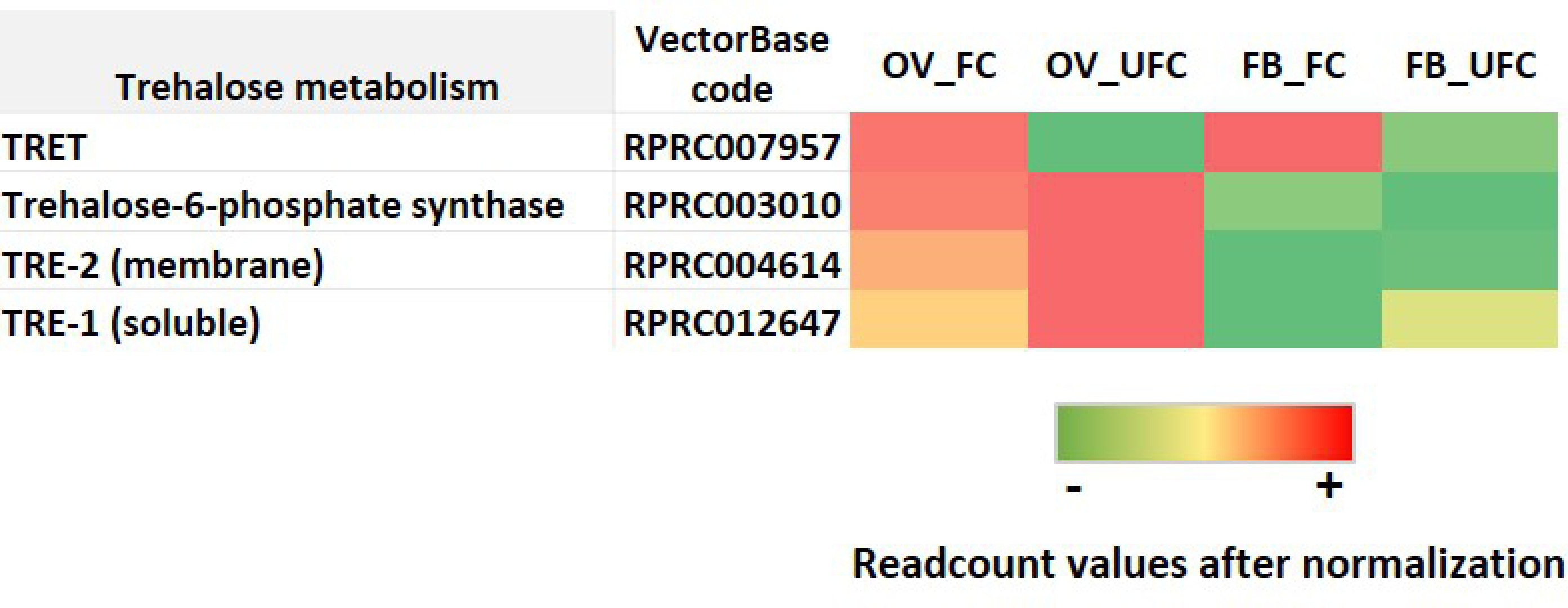
Heat map comparing the mRNA expression levels of molecules involved in trehalose metabolism in fat body and ovaries of female adults in different nutritional condition. The input data is the readcount value from the gene expression level analysis after normalization and is presented by means of a colour scale, in which green/yellow/red represent lowest/moderate/highest expression. DESeq was used to perform the analysis.

**Fig 7.**
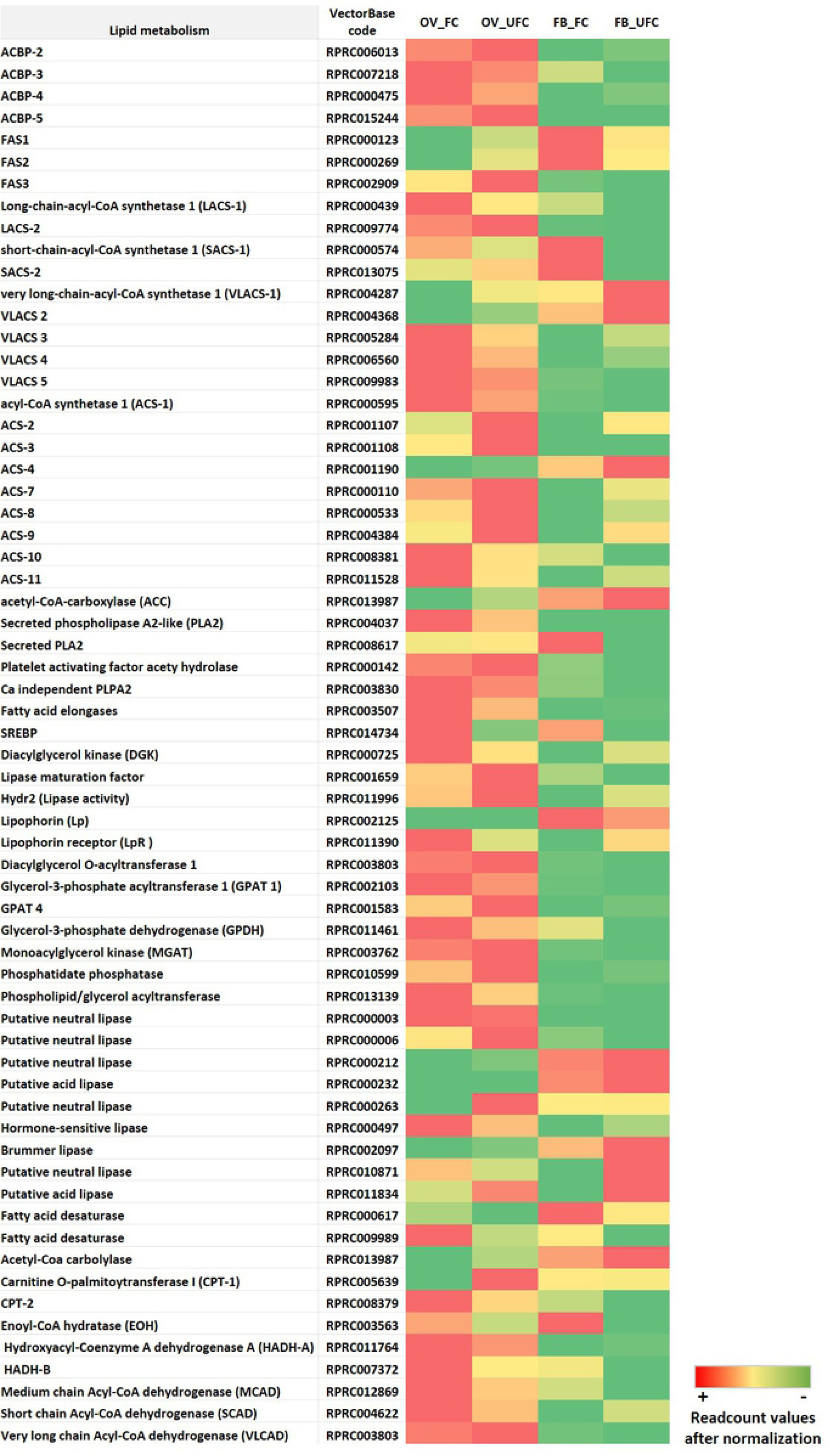
Heat map comparing the mRNA expression levels of molecules involved in lipid metabolism in fat body and ovaries of female adults in different nutritional condition. The input data is the readcount value from the gene expression level analysis after normalization and is presented by means of a colour scale, in which green/yellow/red represent lowest/moderate/highest expression. DESeq was used to do the analysis. DESeq was used to perform the analysis.

### Lipid analysis

In insects the majority of lipid reserves are found in the FB as triacylglycerol (TAG). Lipids are critical to support situations of high metabolic demand, such as vitellogenesis [9]. TAG storage is mainly the result of 2 mechanisms: a) the transfer of dietary fat from the midgut to the FB by lipophorin (Lp), the main lipoprotein of insects, during feeding, and b) the synthesis of lipids from carbohydrates reserves. The participation of lipids in oocytes growth is mainly to supply energy for the developing embryo [23]. As the ability of insect oocytes to obtain fatty acids by *de novo* synthesis is very small, most of the lipids in the oocyte come from the FB via the hemolymph using Lp as transport [66]. In vitellogenesis, lipid accumulation by OVs is associated with a considerable reduction in the lipid content of the FB [9]. However, after a large blood meal, the triatomines must store a vast amount of TAG to support a possible period of fasting. This reality promotes a fine balance between lipid mobilization for egg growth and lipid storage to survive starvation. Here, we demonstrate that there are different types and subtypes of enzymes involved in lipid metabolism, as reported by Gondim et al. [67], and each one seems to have a particular role according to the specific tissue and physiological condition. TAG can be synthesized by 4 different pathways: a) the monoacylglycerol (MG)-pathway; b) the glycerol-3 phosphate (G3P) pathway; (c) degradation of phospholipids; or (d) deacylation of triglyceride catalyzed by lipases [9]. In *R. prolixus*, only the G3P pathway has been reported [67]. This pathway starts with acylation of G3P, catalyzed by G3P acyl transferases (GPAT). Two GPAT, GPAT1 and GPAT4, have been described in *R. prolixus* [68]. Here, we find that the mRNA expression of *GPA1* and *GPA4* is predominantly increased in the OV with respect to the FB and only *GPAT4* is up-regulated in the OV of unfed insects (Fig 7 and S2 Table). We had expected these enzyme to be increased in the FB after a blood meal but it is important to highlight that Alvez-Bezerra et al. [68] indicated that GPAT activity is regulated by a post-translational mechanisms and not at the mRNA levels. However, other transcripts for enzymes involved with the synthesis, elongation and lipid storage, such as *insect microsomal and cytosolic fatty acid synthases (FAS1 and FAS2), lipid elongases and sterol regulatory element-binding protein (SREBP)* are up-regulated in the FB after a blood meal (Fig 7 and S2 Table). These finding coincide with our previous report, where we show that both, “fatty acid biosynthesis” and “fatty acid elongation”, are KEGG pathways enriched in FB_FC [17]. Fatty acid desaturases (FAD) are essentials for *de novo* FA synthesis. In *R. prolixus* we show that 2 transcripts encoding for *FAD* are up-regulated in both FB and OV of fed insects (Fig 7 and S2 Table). These results suggest that after a blood meal, FA synthesis increases and confirms that, besides incorporation of lipids from hemolymph, *de novo* synthesis of FAs by the FB of *R. prolixus* occurs, as was suggested by Pontes et al. [69]. Therefore, FAs stored in tissues could be used to synthesize TAG, phospholipids or be oxidized for ATP production. For any of these pathways, FAs need to be activated and that is the role of acetyl CoA synthetases (ACS). In *R. prolixus*, we find different *ACS* transcripts that encode short-chain ACS, regular ACS, long-chain ACS and very long chain ACS. All these enzymes are present in both the FB and OV, but their expression patterns depend on the nutritional condition (Fig 7 and S2 Table). In general, ACS could be considered more important during the unfed condition, suggesting that β-oxidation is an essential pathway in unfed insects to promote the synthesis of ATP as an energy source (Fig 7 and S2 Table). For FA mobilization, lipases play a critical role to catalyze the hydrolysis of TAG molecules [9]. In this sense, transcripts related to lipid breakdown (*lipases*) or lipid transfer (*lipophorin receptor, LpR*) in general are increased in the FB of unfed insects (Fig 7 and S2 Table). Among others, we also find an increase (not statistically significant) of *Brummer lipase-like* and *Hormone-sensitive lipase-like* mRNA expression in FB_UFC. Interestingly, in *D. melanogaster*, Brummer lipase is induced in the FB during starvation by FoxO-signaling [70]. Recently we showed that Foxo signaling is up-regulated in FB_UFC [17]. Hormone-sensitive lipase is present in the lipid droplet of *D. melanogaster* and is involved in FB lipid mobilization during starvation [71]. Also, it was reported that in insects, the activation of lipolysis is accompanied by hydrolysis of phospholipids from lipid droplets, which suggests that the phospholipase enzyme could be required to allow access of lipases to TAGs contained in the core of the lipid droplets [72]. In this context, Brummer lipase belongs to the calcium-independent phospholipase A2 (iPLA2) family [9]. In *N. lugens*, a deficiency of this enzyme during vitellogenesis impairs lipid mobilization, negatively affecting egg production [73]. The reality that Brummer lipase mRNA expression show only a small increase during UFC (statistically non-significant, S2 Table) could be due to the fact that in *R. prolixus* this enzyme is working the same as in *N. lugens*, being necessary in both nutritional conditions, due to a pleiotropic effect. In addition, the lipase maturation factor 1 is a protein involved in the post-translational maturation of secreted homodimeric lipases [74]. In times of high energy demand, such as starvation, insects use TAG stores via the coordinated action of lipases. In our experiment, *lipase maturation factor* transcript expression is up-regulated in OV_UFC, as is the expression of *Hydr2* (*lipase activity enzyme)*, among other lipases (Fig 7 and S2 Table). These findings are another indication of the fine cross-talk between lipid synthesis and mobilization in both nutritional conditions.

Given the premise that oocytes have a low capacity to synthesize lipids *de novo*, it is surprising to find that *FAS2, FAS3* and *Acetyl CoA decarboxylase* mRNAs, which are lipogenic enzymes involved in *de novo* synthesis of FA, are up-regulated in OV_UFC (Fig 7 and S2 Table). Recently, we reported via KEGG analysis an up-regulation of “fatty acid biosynthesis pathway” in OV_UFC [17]. In mosquitoes, FAS is more highly expressed in diapause-destined females than in non-diapausing individuals [75]. This finding suggests that in *R. prolixus*, FAS could be working to convert carbohydrate reserves to lipid stores for use as an energy source to maintain OVs under optimal physiological conditions for successful reproduction when nutritional conditions are adequate, such as after feeding. Massive endocytosis of YPPs in oocyte and intense VgR, LpR and heavy-chain clathrin synthesis are all energy-dependent processes [76]; for that reason, lipid reserves in pre-vitellogenic oocytes (UFC) could play a critical role in supporting the energetic demands of the growing oocyte at the beginning of vitellogenesis.

In the triatomine, *Panstrongylus megistus*, lipid transfer to the developing oocyte during vitellogenesis is accomplished by endocytosis of Lp (through LpR) and by the classic extracellular lipophorin shuttle mechanism [23]. However, Machado et al. [77] suggested that in *R. prolixus*, endocytosis is not a pathway involved in lipid transfer to oocytes. Conversely, our results demonstrate that *LpR* transcript is up-regulated in OV_FC, probably to maximize lipid delivery to oocytes. Moreover, in mammals, it is known that once lipid levels drop, SREBP induces the expression of many genes involved in lipid synthesis and uptake, including the LDL receptor [78]. It has been reported that SREBP controls lipid uptake and accumulation in oocytes from *D. melanogaster* by regulation of LpR expression [79]. In our data we find up-regulation of *SREBP* mRNA in OV_FC (Fig 7 and S2 Table), suggesting that this transcription factor could be involved in lipid accumulation by the oocytes during vitellogenesis. The difference found in *R. prolixus* [77] could be due to 3 situations: a) an increase of transcript expression not resulting in an increase in the mature protein; b) the difference in the time chosen for the experiments; or c) the use of less sensitive approaches.

Diacylglycerol kinase (DGK) is a family of enzymes that catalyzes the conversion of diacylglycerol (DAG) to phosphatidic acid (PA). We find that *DGK* transcript expression is up-regulated in OV_FC (Fig 7 and S2 Table). PA is a component of the membrane phospholipids and at this stage there is a high demand for membrane synthesis, which is used for oocyte growth and/or for organelles formation, such as yolk granules and lipid droplets. On the other hand, PA affects numerous intracellular signaling pathways, including those regulating cell growth, differentiation, and membrane trafficking. Indeed, PA can bind to mToR and promote ToR signaling [80]. This finding further supports mToR signaling activation after a blood meal in OVs of *R. prolixus* [17]. Also in insects, the requirement of an acyl-CoA synthetase long chain (ACSL1) for oviposition and egg viability has been reported [81]. In our work, we find up-regulation of *ACSL1* mRNA in OVs during FC (Fig 7 and S2 Table). Acyl-CoA-binding protein (ACBP) are small proteins that binds acyl-CoA esters with very high affinity to protect them from hydrolysis.

Although *ACPB-2, ACPB-3, ACPB-4* and *ACBP-5* transcripts are present in both tissues, only *ACPB-3* is up-regulated in FB_FC meanwhile *ACPB-4* is up-regulated in OV_FC (Fig 7 and S2 Table), indicating that the involvement in TAG mobilization by ACPB is specific and unique, depending on the tissue and physiological condition.

### Neuropeptides and neurohormonal signaling, and serotonin

A variety of neuropeptides and neurohormones have been identified in the CNS of *R. prolixus* [82]. FB and OV development and function are largely regulated by several hormonal and nutritional signals, i.e. ILP/ToR signaling [17]. Our transcriptome analysis showed no significant change in mRNA expression after blood intake in CNS. However, we made a deep analysis in CNS, FB and OV to explore the relative expression of transcripts related to hormonal signaling in both nutritional conditions. Here, we discuss neuropeptides, in addition to the amine serotonin, and their receptors, which show high expression in some of the tissues analyzed (for more details see S2 Table). All neuropeptides are synthesized as part of a larger precursor molecule. The selective processing of those precursors determines which peptides are finally released by the specific cells [83]. Here, we find 7 enzymes involved in neuropeptide processing and all of them are expressed in the CNS, FB and OV in both nutritional conditions (Fig 8A and S1 Table). The results support the contribution of FB and OV for neuropeptide production in both nutritional condition.

**Fig 8.**
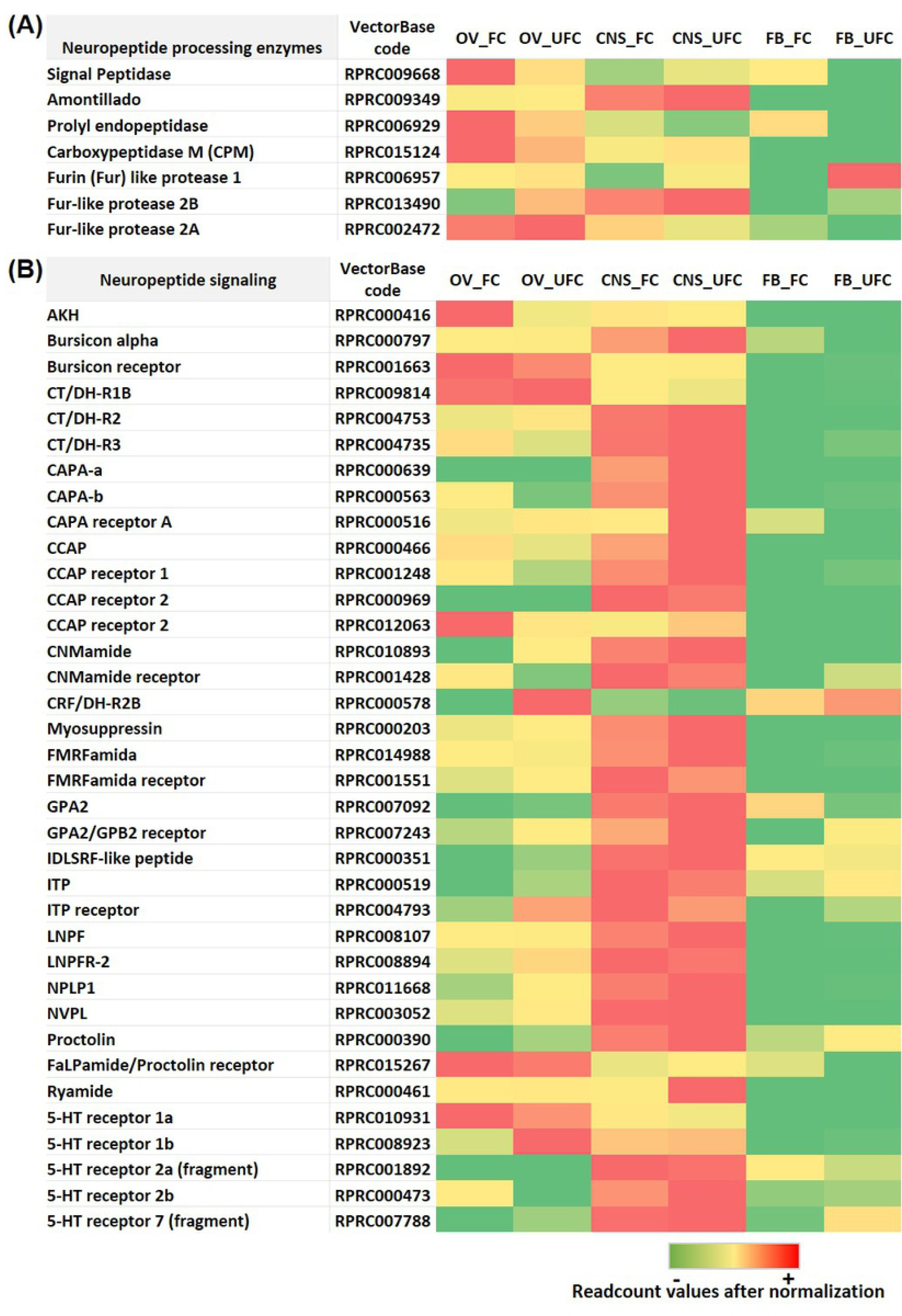
Heat map comparing the mRNA expression levels of molecules involved in neuropeptide processing enzymes (A) and neuropeptide signaling (B) in fat body, ovaries and CNS of female adults in different nutritional condition. The input data is the readcount value from the gene expression level analysis after normalization and is presented by means of a colour scale, in which green/yellow/red represent lowest/moderate/highest expression. DESeq was used to perform the analysis.

The presence of the AKH precursor and receptor [84] in OVs suggests a role in egg production and/or egg-laying behaviour as has been shown in other insects [85], possibly by an autocrine pathway. Here, we find that *AKH* transcript expression is detected in CNS but is up-regulated in OV_FC (Fig 8B and S2 Table). In insects, buriscon is a heterodimeric glycoprotein hormone which plays a key role in melanization and cuticle hardening during development of insects [86]. Recently, a novel function of bursicon was reported in the stimulation of Vg expression in the black tiger shrimp, *Penaeus monodon* [87]. In *R. prolixus*, we find higher expression of the *bursicon receptor* in OVs with respect to the CNS and FB (Fig 8B and S2 Table), suggesting a novel role for this hormone in reproductive physiology in an insect. Human genome screening reveals the presence of another glycoprotein hormone, consisting of the novel alpha (GPA2) and beta (GPB5) subunits (GPA2/GPB5) [88]. In *A. aegypti*, GPA2/GPB5 signaling has been implicated in controlling ionic balance [89]. In addition, this signaling pathway could play a role in spermatogenesis and oogenesis in male and female mosquitoes, respectively [90]. We find an up-regulation of *GPA2/GPB5 receptor* mRNA expression in OV and FB during UFC, suggesting an involvement of this signaling pathway in the stage prior to vitellogenesis (Fig 8B and S2 Table). Also, in rats, it has been reported that GPA2/GPB5 in the ovary may act as a paracrine regulator in reproductive processes [91]. Our results show up-regulation of *GPA2* mRNA in OV_UFC and conversely, up-regulation of this transcript in FB_FC (Fig 8B and S2 Table). Future experiments will determine the involvement of this new signaling pathway in insects and its interplay with reproductive processes. Calcitonin-like diuretic hormones (CT/DHs) are related to the mammalian calcitonin and calcitonin gene-related peptide hormonal system. Here, in addition to the expression in CNS, we show a high mRNA expression level of *CT/DH-R*s in OV with moderate levels in the FB (Fig 8B and S2 Table). Previously, in *R. prolixus*, it was suggested that CT/DH-Rs signaling may have a critical, but unknown, role in reproductive physiology [92].

*R. prolixus* genome has two paralogue genes encoding capability (CAPA) peptides, named *RhoprCAPA-α* and *RhoprCAPA-β* [93-94]. These genes are mainly expressed in the CNS, supporting our transcriptome results (Fig 8B and S2 Table). RhoprCAPA-α expression was also detected in testes from 5th instar nymphs but not from adults, suggesting a role in the maturation of male gonads [93]. Here, we find *CAPA-β* transcript expression is up-regulated in OV_FC. Future experiments using gene silencing strategies will be performed to analyse the possible involvement of CAPA-β peptides on oocyte maturation or egg formation.

Pleiotropic effects of crustacean cardioactive peptide (CCAP) in insects and crustaceans have been described. Previously, it was reported that CCAP is involved in the fertilization process in *L. migratoria* since it increases the basal tonus and frequency of spontaneous spermathecal contractions [95]. Our results show an up-regulation of *CCAP* mRNA expression in OV_FC (Fig 8B and S2 Table), suggesting an autocrine regulation but future experiments are required to determine the specific involvement of this signaling in *R. prolixus* reproduction.

Ion transport peptides (ITPs) in locusts (*Schistocerca gregaria* and *L. migratoria*) were identified based on their antidiuretic activity on the ileum [96-97]. Later, in *T. castaneum* it was suggested that ITP signaling participates in ovarian maturation and female fecundity regulation [98]. However, its specific role in reproductive physiology in *R. prolixus* has not yet been reported. Here, we found an up-regulation of *ITP receptor* mRNA in both FB and OV from fed insects (Fig 8B and S2 Table).

In insects, long neuropeptide F (LNPF) has been reported as a main player in feeding behaviour, metabolism and stress responses [99]. Previously, in *R. prolixus*, it was reported that pre-follicular cells within the germarium express the NPF receptor, as do cells located between developing oocytes [100]. Taking into account both findings, our results suggest that LNPF signaling could have a critical role in oocyte maturation more than in egg production, since we find an up-regulation of *LNPF receptor* mRNA in OV_UFC with respect to FC (Fig 8B and S2 Table).

Neuropeptide-like precursor 1 (NPLP1) was first identified in *D. melanogaster*. In *R. prolixus* NPLP1 peptides are involved in the feeding response, providing the first clues in the elucidation of their function [21]. We find an up-regulation of *NPLP1* transcript expression in OV_UFC (Fig 8B and S2 Table). The physiological role of NLPL1 signaling in reproduction is currently unknown.

By quantitative peptidomic assays, it was reported that in *R. prolixus*, NVP-like (NVPL) signaling is involved in the regulation of rapid events, such as diuresis/antidiuresis, and in delayed events such as mating and reproduction [21]. In our transcriptome analysis, we show an up-regulation of *NVPL* mRNA in OV_UFC (Fig 8B and S2 Table). Gene silencing techniques could be implemented to evaluate the role of this peptide in reproduction.

Myosuppressin is a neuropeptide only found in insects and crustaceans. It has been demonstrated to have anti-feeding activity and to inhibit gut and oviduct contraction and neuropeptide secretion [101]. In the Australian crayfish *Cherax quadricarinatus*, myosuppressin was detected in ovaries from mature females, suggesting a potential link between myosuppressin and reproduction [102]. Here, we also report *myosuppressin* mRNA expression in OVs of *R. prolixus* (Fig 8B and S2 Table).

A corticotropin-releasing factor-like peptide acts as a diuretic hormone in *R. prolixus* (Rhopr-CRF/DH) [103]; however, its distribution throughout the CNS and the expression of its receptor in feeding-related tissues as well as the female reproductive system suggests a multifaceted role for the neuropeptide. Adult female *R. prolixus*, injected with Rhopr-CRF/DH produce and lay significantly fewer eggs [104]. In addition, in locusts, CRF/DH inhibits oocyte growth and reduces ecdysteroid levels [105]. Here, we find an up-regulation of *CRF/DH receptor* mRNA in OV and FB from unfed insects (Fig 8B and S2 Table), where vitellogenesis is inhibited, supporting its effects as a negative regulator of reproduction.

By bioinformatic predictions, Ons et al. [105] showed for the first time the existence of RYamide in *R. prolixus*. However, the functions of this signaling in insects is currently unclear. We find a high expression of *RYamide* mRNA in OVs during both nutritional condition (Fig 8B and S2 Table).

Proctolin was the first insect neuropeptide to be sequenced and synthesized and is found in a variety of arthropods, including *R. prolixus* [107], where it plays a myostimulatory role on anterior midgut, hindgut, heart, and reproductive tissue [108]. In the cockroach *Blaberus craniifer*, nanomolar quantities of proctolin induce Vg uptake [109]. Here, we find for first time a high expression of *proctolin receptor* mRNA in OVs, encouraging further studies to analyze the role of this signaling in the reproductive organs (Fig 8B and S2 Table).

Serotonin (5-hydroxytryptamine or 5-HT) is an ancient monoamine neurotransmitter/neurohormone. 5-HT receptors are classified based on sequence similarities with their counterparts in vertebrates [110]. In *R. prolixus*, we find that mRNA expression to *5-HT receptors* is higher in the CNS but also expressed in the OV and FB (Fig 8B and S2 Table). In mosquitos, 5-HT2B was reported to be a critical player in the fat body-specific serotonin signaling system, governing antagonistic ILP actions [111]. It would be interesting to analyse 5-HTs functional role in reproductive tissues of *R. prolixus*.

The transcriptome data highlights directions for future research in examining the role of particular neuropeptides/amines on specific responses to processes such as ovarian maturation or egg formation, extending the temporal range of transcript/protein expression of these neuropeptides/amines capitalizing on gene silencing assays.

### A brief analysis of genes related to immunity

The overall achievement of insects in maintaining a stable population of individuals is due, in part, to their ability to recognize pathogens and eliminate them successfully using the immune system. The immunity of insects comprises multiple elements that work in concert and, in general, includes physical barriers as well as innate immune responses, which lead to a combination of cellular and humoral immunity [112]. In recent years, it has been shown that reproduction and immunity can be mutually constraining since both responses are energetically costly, and therefore need to be traded off. In this context, increased reproductive activity reduces constitutive and induced immunity across a diversity of female insects [113]. However, metabolic changes that occur after the acquisition of a blood meal, include the induction of oxidative stress [114]. Increased metabolic activity during the process of blood digestion has been shown to alter levels of different detoxification enzymes in mosquito, which are the same as these implicated in insecticide detoxification; indeed blood feeding status in mosquitos confers increased tolerance to insecticides [115]. Thus, it is clear that the immune system is working in both nutritional conditions, before a blood meal, due to the stress that is generated by starvation, and after a blood meal, due to the potential toxicity of the molecules ingested with the blood. Along with all the roles described above for FB in reproduction, the FB also responds to microbial infection. One important humoral response is the production of inducible antimicrobial peptides (AMPs), which are rapidly synthesized after microorganism invasion [116]. In *D. melanogaster*, the Toll pathway (activated by fungi and gram-positive bacteria) and the Imd pathway (activated by gram-negative bacteria) lead to the synthesis of AMPs, not only by a pathogenic challenge, but also by aging, circadian rhythms, and mating [118-120]. Interestingly, in *R. prolixus*, we find an up-regulation of AMPs in OV_FC (Fig 9A and S3 Table), suggesting a role of the vitellogenic oocytes in humoral immunity, an event that has not yet been studied in insects. In addition, we find different mRNAs involved with both Toll and Imd pathways which are up- and down-regulated in FB and OV, without revealing a characteristic expression pattern in any of the nutritional conditions analyzed (Fig 9B, C and S3 Table). This finding clearly suggests that the immune system is responding to both stimuli: to detoxification of compounds which enter with blood intake and/or to avoid tissue damage due to stress caused by lack of food. In addition, Foxo transcriptional factor could promote activation of the stress-responsive Jun-N-terminal kinase (JNK) pathway, which antagonizes ILP signaling in *D. melanogaster*, causing nuclear localization of FoxO and inducing its targets, including growth control and stress defense genes [120]. Recently, we demonstrated that in unfed females, FoxO factor is translocated to the nucleus, stimulating the insulin-sensitive pathway and modulating longevity signaling in *R. prolixus* [17]. In the current work, we find up-regulation of most of the genes involved with JNK signaling, mainly in OV_UFC (Fig 9D and S3 Table) possibly to overcome effects of stress and low nutrition.

**Fig 9.**
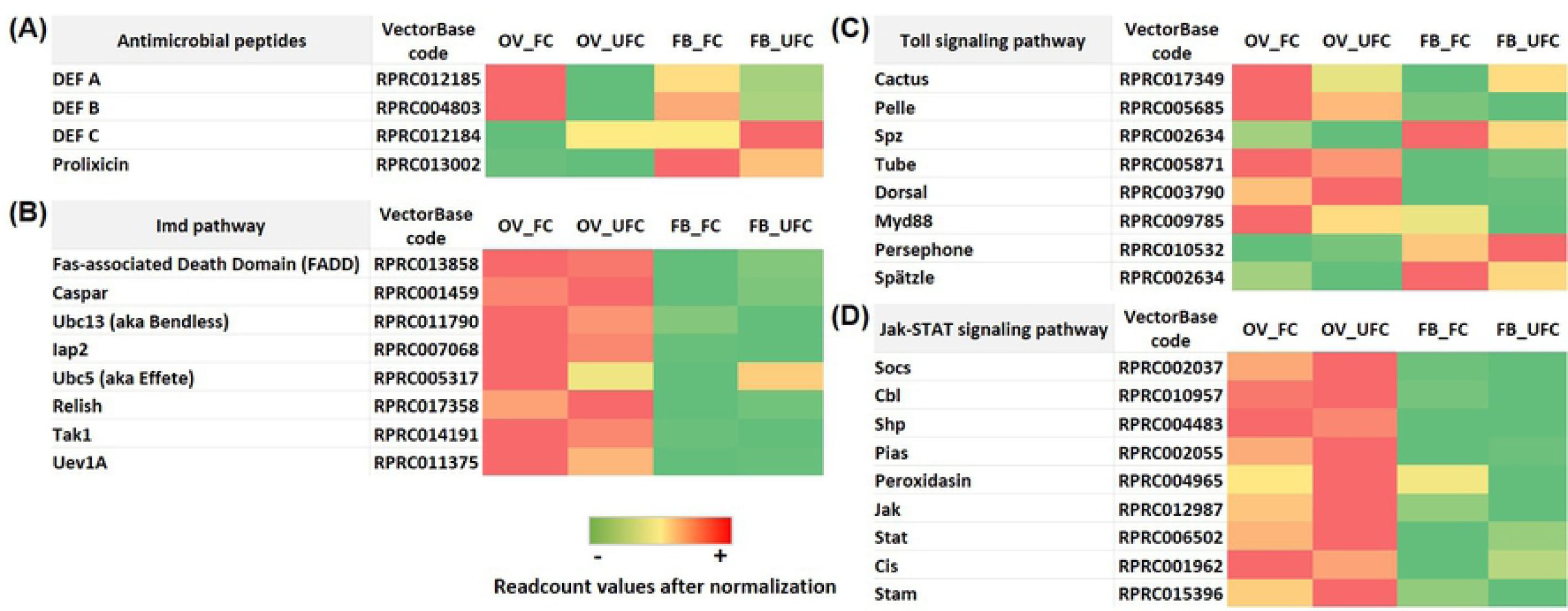
Heat map comparing the mRNA expression levels of molecules involved with Antimicrobial peptides (A), Imd pathway (B), Toll signaling pathway (C) and Jak-STAT signaling pathway (D) in fat body and ovaries of female adults in different nutritional condition. The input data is the readcount value from the gene expression level analysis after normalization and is presented by means of a colour scale, in which green/yellow/red represent lowest/moderate/highest expression. DESeq was used to perform the analysis.

In *R. prolixus*, Duox is the enzyme that generates H_2_O_2_ in ovarian follicles is used as a fuel for hardening of eggshell proteins, a process essential for the acquisition of resistance to water loss [121]. In accordance with those finding, we show an up-regulation of *Duox* mRNA expression in OV_FC (Fig 10A and S3 Table). In addition, melanization and the production of nitric oxide (NO) and reactive oxygen species (ROS) are effector mechanisms also activated as a first line of defense. Upon infection, pattern recognition receptors activate downstream serine protease cascades that culminate in the activation of prophenoloxidase (PPO), a precursor activated by proteolytic cascades to phenoloxidase for *de novo* synthesis of melanin. NO is highly toxic for a wide variety of pathogens and is produced by nitric oxide synthase (NOS). ROS are produced by conserved nicotinamide adenine dinucleotide phosphate (NADPH) enzymes. Here, we find up-regulation of *PPO, NOS* mRNA levels and increases of *NADPH* transcript in OV_UFC (Fig 10A and S3 Table). Also, it is interesting to see that enzymes which performs as antioxidant elements, such catalases, thioredoxin peroxidases and glutathione peroxidase have their mRNA levels up-regulated in OV and FB from fed insects (Fig 10B and S3 Table). RNA interference (RNAi) is triggered by endogenous or invading double-stranded RNAs (dsRNAs) that arise from hairpin structures, transposable elements, or virus infections [122]. In *R. prolixus* we show that in general, there is an up-regulation of mRNA molecules involved with RNAi signaling in OV_UFC (Fig 10C and S3 Table). These results suggest immunological signaling in OV of unfed insects, possibly to prevent damage during unfavorable metabolic conditions.

**Fig 10.**
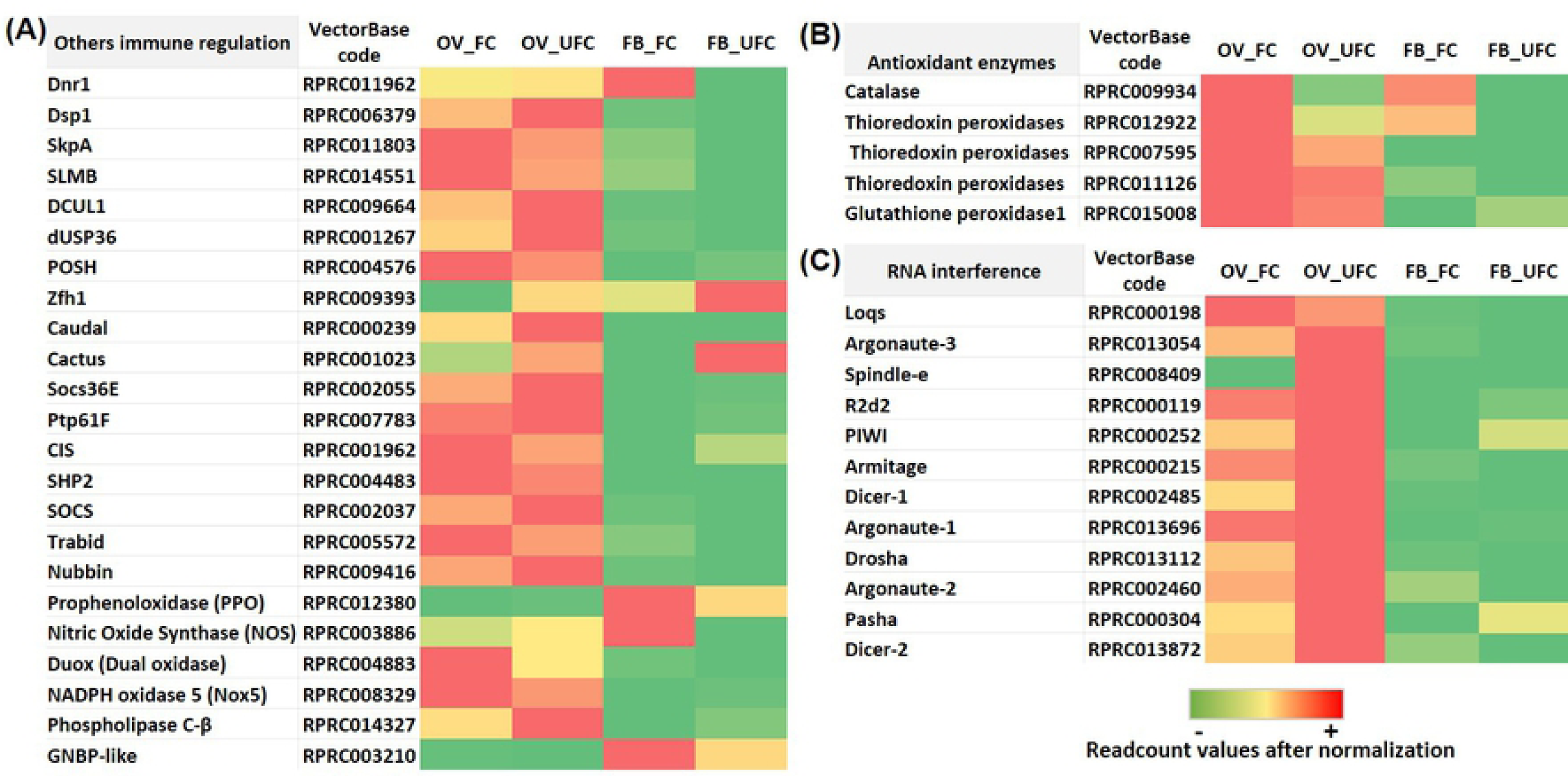
Heat map comparing the mRNA expression levels of various immune regulators (A), antioxidant enzymes (B) and RNA interference signaling (C) in fat body and ovaries of female adults in different nutritional condition. The input data is the readcount value from the gene expression level analysis after normalization and is presented by means of a colour scale, in which green/yellow/red represent lowest/moderate/highest expression. DESeq was used to perform the analysis.

Overall, the information on immunity in hemipterans, including Triatominae vectors remains incomplete and fractionated [123]. The data presented here on immunity and reproduction in triatomine females encouraging the development of future studies to shed light on the relative contribution of the immune system in successful reproductive events.

## Conclusions

We present here a comprehensive analysis of mRNA expression of components of biological processes related with feeding and reproduction. Broadly, using high-throughput sequencing and a comparative expression analysis we find that a blood meal taken by *R. prolixus* females has both unique and interacting effects on CNS, OV and FB gene expression, with patterns of mRNA levels that are consistent with different needs according to the nutritional condition. Of particular interest, we show the cross-talk between reproduction and a) lipid, trehalose and protein metabolism, b) neuropeptide and neurohormonal signaling, and c) the immune system. Overall, our findings provide an invaluable molecular resource for future novel investigations on different tissues related with successful reproductive events, before and after the appropriated stimuli (blood meal). Our data opens up avenues of translational research that could generate novel strategies of vector population control. This includes the identification of specific genes for use in symbiont-mediated RNAi; a powerful technology which provides the potential for biocontrol against tropical disease vectors. In *R. prolixus*, the ability to constitutively deliver dsRNA by supplying with recombinant symbiotic bacteria generated against specific target genes involved in the reproductive success (Vg), have already been tested in laboratory trials and is effective in dramatically reducing the fitness of *R. prolixus* [18].

## Acknowledgement

This research was supported by NSERC Discovery grants to A.B.L. and I.O.

## Author Contributions

J.L., A.B.L. and I.O. designed the experiments and mapped out the manuscript. J.L. performed the experiments, wrote the manuscript and prepared all the figures. A.B.L. and I.O. reviewed and contributed to the writing of the manuscript.

## Competing Interests

The authors declare no competing interests.

## Supporting information

**S1 Fig. Correlation of Log**_**2**_**Fold Change values in fat body (FB) and ovaries (OV) obtained by RNAseq and RT-qPCR data from 7 genes.** Primers used are displayed in S1 Table. The correlation coefficient between RNAseq (y-axis) and RT-qPCR (x-axis) data (log2 fold-change) analyzed by the Pearson test were 0.9311 (a) and 0.9109 (b), with a statistical significance p<0.01.

**S1 Table. Primers used by RT-qPCR assays.**

**S2 Table. Details of the mRNA expression for Figs 4 and 8**. Columns are: the gene name we are assigning; VectorBase code – the official gene number in the RproC3 genome assembly; OV_FC, OV_UFC, FB_FC, FB_UFC, CNS_UFC and CNS_FC show the readcount after normalization. Log_2_FoldChange: log*2* (fed condition/unfed condition); *p-*adj: *p*value after normalization (the smaller the *p-*adj, the more significant the difference). Excel cell highlights in green: up-regulation in fed condition; excel cell highlights in orange: up-regulation in unfed condition. CNS_FC, central nervous system post-feeding (FC, fed condition); CNS_UFC, central nervous system before of a blood meal (UFC, unfed condition); FB_FC, fat body in FC; FB_UFC, fat body in UFC; OV_FC, ovary in FC; OV_UFC, ovary in UFC.

**S3 Table. Details of the mRNA expression of Figs 9 and 10.** Columns are: the gene name we are assigning; VectorBase code – the official gene number in the RproC3 genome assembly; OV_FC, OV_UFC, FB_FC and FB_UFC are the readcount after normalization. Log_2_FoldChange: log_2_ (fed condition/unfed condition); *p-*adj: *p*value after normalization (the smaller the *p-*adj, the more significant the difference). Excel cell highlights in green: up-regulation in fed condition; excel cell highlights in orange: up-regulation in unfed condition.

